# Insights into the ligand-free structure of cyclic diguanosine monophosphate I riboswitch of *Vibrio cholerae* using molecular dynamics simulation

**DOI:** 10.1101/871194

**Authors:** Priyanka Kumari, Anup Som

## Abstract

Riboswitches are key cis regulatory elements present at 5’ UTRs of mRNAs. They play a critical role in gene expression regulation at transcriptional and translational level by binding selectively to specific ligands followed by conformational changes. Ligands bind to the aptamer of riboswitches and their complex structures have been solved, but ligand-free riboswitches structures are not available which is important to understand specific ligand binding mechanism. In this paper, an all atom 150 nano-second (ns) molecular dynamics (MD) simulations of cyclic diguanosine monophosphate (c-di-GMP I) riboswitch aptamer domain from *Vibrio cholerae* were carried out to study ligand-free c-di-GMP I riboswitch aptamer structure and the binding mechanism. The Principle component analysis, cross correlation dynamics analysis and trajectory analyses revealed that the ligand-free structure has stable conformation with folded P2, P3 and an open P1 helix which opens the ligand binding helix-join-helix while the ligand-bound structure shows less deviation and remains as closed structure compared to the ligand-free structure. The junction residues significantly showed anti-correlated motions with each other elucidating the open conformation of the ligand-free aptamer of riboswitch. The identified key residues involved in binding are A18, G20, C46, A47 and C92.

**Highlights:**

- The c-di-GMP I riboswitch regulates the essential genes involved in the virulence of human bacterial pathogen *V. Cholera*.
- A 150 ns molecular dynamics run was performed to find a ligand-free stable structure of c-di-GMP I riboswitch aptamer.
- The trajectory analysis resulted in stable conformation of ligand-free structure with folded P2, slightly open P3 and an unwind P1 helix.
- The atomic level analyses through cross correlation dynamics and RMSF values showed the opening of catalytic pocket and unwinding P1 helix.
- The identified key residues involved in binding are A18, G20, C46, A47 and C92 at the catalytic pocket.

## Introduction

Riboswitches are cis-acting, gene regulatory structural domains usually present at the 5’ UTR region of mRNA which selectively recognizes specific small ligands/metabolites (Barrick and Breaker, 2007; Guil and Esteller, 2015). The structure of riboswitch consists of two regions; aptamer and expression platform. The aptamer binds to specific ligand and regulates the adjacent gene/genes through conformational changes induced at expression platform during transcription, translation and alternate splicing (McCown 2017). Thus riboswitches are critical to organisms’ life processes and represent a new class of RNA antibiotic drug target for infectious diseases especially bacterial diseases (Blount and Breaker, 2006; Kumari and Som, 2019; Mehdizadeh et al., 2016; Rekand, 2017; Chan et al., 2017). Riboswitches regulate gene expression in bacteria, archeae, fungi and some eukaryotes.

On the basis of the type of ligand bound to a riboswitch, more than 40 classes of riboswitches have been discovered (McCown 2017). One such class of riboswitch is Cyclic-di-GMP I ribowsitch which senses cyclic diguanosine monophosphate (c-di-GMP/cyclic-di-GMP) metabolite through its aptameric portion. It folds into secondary and tertiary configurations forming a stable three dimensional structure. The cyclic-di-GMP I ribowsitch controls the expression of genes involved in the metabolism of cyclic-di-GMP which play crucial role in bacterial survival (Sudarsan et al., 2008). The cyclic-di-GMP I is a secondary messenger involved in the mechanisms such as virulence, mobility, quorum sensing and biofilm formation. McKee et al. (2018) showed that the c-di-GMP regulates virulence factor genes via riboswitches in *C. difficile*. In a recent study, Tamayo (2019) reported the role of c-di-GMP riboswitches in controlling bacterial pathogenesis mechanisms. They are present in wide variety of bacteria specifically the pathogenic stains of proteobacteria (Kumari and Som, 2019). Therefore, targeting these riboswitches might prove beneficial as they had been proposed as therapeutic drug target (Rekand, 2017). The conformational transition occurring in bound and unbound riboswitch with its metabolite governs the expression of genes adjacent to them, but in data repositories only information about riboswitch with ligand complex is available. Thus it is necessary to get information about the initial ligand-free structure to study the mechanism of aptamer-ligand binding and gene regulation. Molecular Dynamics (MD) studies has been proved valuable to gain insight into the structural and functional aspects of biomolecules and had been used earlier to solve similar problems, for example: Gong et al (2014) used MD simulations to study ligand binding to PreQ1 riboswitch using trajectory analyses such as root mean square deviation (RMSD), root mean square fluctuations (RMSF) and solvent accessible surface area (SASA) calculations. A similar work was carried out by Sharma et al (2009) for add A-riboswitch to show the role of secondary structural orientation and motions for binding mechanism. A recent approach is the use of replica exchange MD simulation for enhanced sampling as one of the drawbacks of MD simulation is insufficient sampling due to time and computational constrains to reach global minima (Xue et al., 2015). Further, in a study Li et al (2018) showed the use of MD studies to analyse the binding of natural and inhibitor ligand to 3’-3’-cGAMP riboswitch. The utility of MD simulations in understanding the RNA structural dynamics is thoroughly discussed in a review article by Sponer et al (2018). Thus, in this paper MD simulation based study was conducted to predict the ligand-free cyclic-di-GMP I aptameric ribowsitch and the changes in the aptameric riboswitch with ligand binding. In order to characterize the ligand-free structure, understanding the secondary and tertiary structure of the cyclic-di-GMP I riboswitch aptamer is prerequisite.

### Structure of c-di-GMP I riboswitch aptamer

To study the conformational changes and mechanism, detailed knowledge of secondary and tertiary structure is important. The secondary structure of cyclic-di-GMP I riboswitch 6 aptamer shows three helices P1, P2 and P3 forming a three way junction representing junction loops J1/2, J2/3 and J1/3 respectively (Figure 1). The helix P1 represents nucleotides from G9-C13 and G94-C97, the helix P2 from C22-G45 and the helix P3 from C51-A91. The c-di-GMP I riboswitch aptamer binding pocket is composed of residues from the junction of the three helices (P1, P2, P3) known as junction loops J1/2 J1/3 and J2/3. The nucleic bases present in the ligand binding cavity are represented in Table 1. This might be the area of interest to analyse the dynamics of the molecules.

**Table 1:**
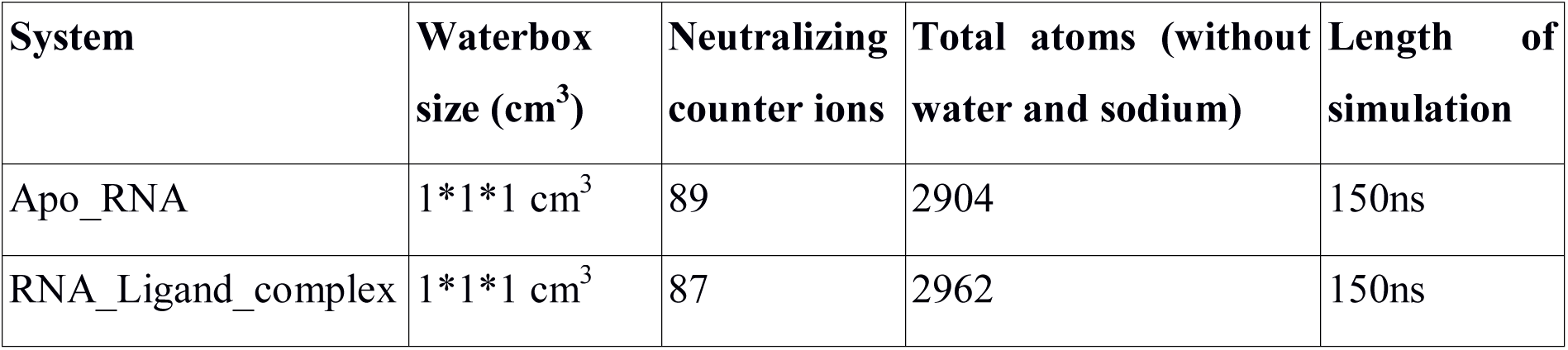
Details of systems preparation for MD simulations.

| System | Waterbox size (cm<sup>3</sup>) | Neutralizing counter ions | Total atoms (without water and sodium) | Length of simulation |
| --- | --- | --- | --- | --- |
| Apo_RNA | 1*1*1 cm<sup>3</sup> | 89 | 2904 | 150ns |
| RNA_Ligand_complex | 1*1*1 cm<sup>3</sup> | 87 | 2962 | 150ns |

**Figure 1:**
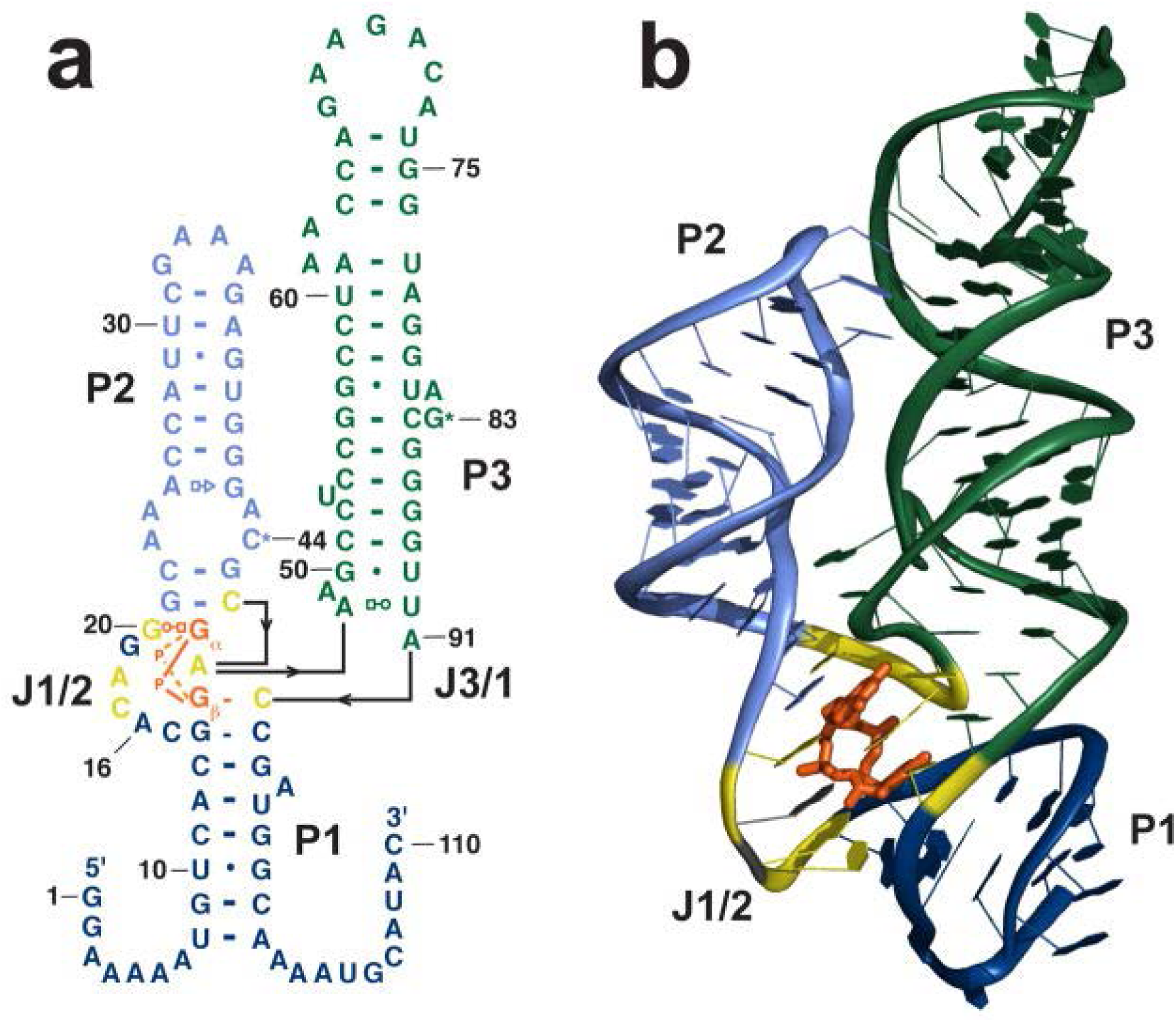
Secondary and tertiary structures of c-di-GMP I riboswitch. (**A**) Represents the secondary structure and (**B)** represents the tertiary structure. Helices P1, P2 and P3 are represented by navy blue, blue and green colour and the junction loops J1/2, J2/3 and J1/3 of the binding pocket is represented in yellow colour. The ligand C2E is shown in orange colour.

## Materials and Methods

An all atom MD simulations of 150 ns were used for this study. The systems along with the details of the software and algorithms used are as follows:

### Initial structure used

The X-ray crystal structure of c-di-GMP I riboswitch aptamer used for the study was retrieved from protein data bank (PDB) database [PDB id – 3IRW] (Smith et al., 2009). The resolution of the crystal structure is 2.7 Å that consist of two chains one of protein and other of RNA. The RNA chain has 90 nucleotides. This structure was used as template to derive two systems for the MD simulations.

### System preparation from initial structure

For this study two systems were used: Apo_RNA system (ligand-free) and RNA_ligand_complex system (ligand-bound). Apo_RNA system was prepared by removing first the protein chain then the hetero-atoms such as C2E ligand, water molecules, magnesium ions (Mg^+2^) and IRI molecules using Chimera tool (Pettersen et al. 2004). For the RNA_ligand_complex system, the protein chain, water and IRI molecules were removed from the crystal structure. The systems were placed at the centre of cubic boxes, then each box was solvated using explicit water system TIP3P (Jorgensen et al., 1983) The partial charges of the systems were neutralized by adding monovalent ions to them (details given in Table 1). Due to addition of water and ions to the system, steric hindrance might manifest between these molecules. Thus two systems were energy minimized using steepest descent minimization algorithm, a method of optimization (Freund, 2004) with a maximum force cut off value <1000 Kcal/mole/nm with periodic boundary conditions for 50000 steps. The energy minimization is done to remove bad contacts and optimize the structure.

### MD simulation

All MD simulations were carried out by using GROningen MAchine for Chemical Simulations (GROMACS) 5.1.2 package (Abraham et al., 2015) with AMBER99SB-ildn force field. The force field for the ligand was generated using AnteChamber PYthon Parser interfacE (ACPYPE) server (da Silva and Vranken, 2012). Before the production dynamics the energy minimized system were equilibrated using NVT [number of atoms, volume and temperature] and NPT [number of atoms, pressure and temperature] ensambles. During NVT equilibration for 100 picoseconds (ps), the system is heated from 0 to 300 K and the temperature is maintained at a constant 300 K using the modified Berendsen thermostat (Berendsen et al., 1984). This equilibration assigned initial velocities to the atoms of the system from Maxwell distribution. The next equilibration step is NPT equilibration, run for 100 ps dynamics to maintain a constant pressure of the system by Parinello-Rahman barostat (Parinello and Rahman, 1981). It also maintains the homogeneous density of the system. Thus the system was maintained at room temperature, atmospheric pressure and homogenous density. The Linear Constraint Solver (LINCS) algorithm is used for short range electrostatic and van der Waals interactions and the cut off used is 1.0 nm. For long range electrostatics Particle Mesh Ewald (PME) method was incorporated. The MD simulations were run based on leap-frog algorithm (Van Gunsteren and Berendsen, 1988). The time step used for MD simulation was 2 femtoseconds (fs) and the total time for MD run was 150 ns.

### Trajectory and structure analysis

The trajectories were analysed using VMD (Visual Molecular Dynamics) visualization and analysis tools. VMD was used to visualise all the trajectory movements for the simulation time and extraction of co-ordinates for particular time frame and the average RMSDs. The structure comparison and analyses were done using PyMOL (Schrodinger, 2019) and CHIMERA visualization tools (Pettersen et al. 2004). The RMSD, RMSF, SASA etc. graphs of trajectories were obtained using terminal-shell commands and plotted using XMGRACE 2D plotting tool (Turner, 2005).

### Principle component analysis and correlation dynamics analysis

Principle Component Analysis (PCA) is used to study the relationship of various conformations sampled during the MD production run and helps to analyze the dominant trajectories from the corresponding simulations (David and Jacobs, 2014). PCA for the ligand-free system was generated using Bio3D library of R package (Grant et al. 2006). The covariance matrices were constructed for the backbone atoms for each frame of the 150 ns long dynamics. The graphs of first three principle components PC1, PC2 and PC3 were generated as the first few PCs covers most significant motions. The dynamical changes of the system over the time can be observed by constructing dynamic cross correlation (DCC) graph. This graph elucidates insights into the correlated motions between the different regions of the biomolecule and was generated using Bio3D. The extent of the correlated motion was calculated by correlation coefficient C(i,j) between the i^th^ and j^th^ atoms which is represented by the following equation

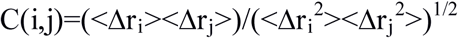

where Δr_i_ and Δr_j_ is the displacement of i^th^ and j^th^ atom from mean position and <> symbol represents the time average.

### Binding pocket analysis

To understand the binding mechanism, it is important to identify the area where the ligand binds, the residues corresponding to it, the hydrogen bonds formed and the acceptor and donor hydrogen bond atoms of the residues. We used chimera tool to determine the binding pocket and other analysis. The residues around the ligand within 5 Å zone is taken as binding pocket residues and the hydrogen bonds formed were retrieved using Hbond analysis tool.

## Results and Discussion

### Stable ligand-free aptamer structure

To identify the possible stable ligand-free aptamer structure of c-di-GMP I riboswitch aptamer which binds to c-di-GMP molecule naturally, the trajectory studies such as RMSD and RMSF parameters were used. RMSD is used as a measure the conformational stability of a structure during the simulation. To evaluate the conformational changes in the ligand-free and ligand-bound states of the riboswitch, the comparative plots of RMSD versus time (Figure 2) for global (complete RNA) and local (P3 helix and binding pocket region) RMSDs were obtained for 150ns MD run. The RMSD graph for the RNA system was generated using the starting minimised structure as reference structure. The RMSD graphs showed atomic-resolution details of their structural dynamics and elucidated several features of ligand-free and ligand-bound structures. Both the ligand-free and ligand-bound structures being flexible and dynamic show variations in RMSD values, the average RMSD for ligand-free structure is 6.44 Å and for ligand-bound structure is 5.90 Å (details given in Table 2). As expected the average RMSD value for ligand-bound structure is lower than ligand-free structure because ligand-bound structure is stable and the removal of ligand from the bound structure renders instability in the ligand-free structure. The structures show stable configuration after 80 ns. Before 80 ns unusual fluctuations were observed for ligand-bound structure. The riboswitch when bind to its native ligand shows lower RMSD values as compared to apo_RNA stating that the stability of the RNA increases when bound to its native ligand. The fluctuations in case of RNA bound to ligand and Mg^2+^ ion showed equilibrium state over the simulation time individually but its RMSD values gets higher than ligand-free structure for 25 to 40 ns. Among the helices, P2 and P3 exhibited stability with lower fluctuations for both ligand-free and ligand-bound simulations (Figure 2, Table 2). However, the helix P1 contributed to the conformational change as the RMSDs of ligand-free structure (10.8 Å) were higher and stable as compared to ligand-bound structure (8.87 Å). The fluctuations at the P1 helix also contributed to the opening of the binding site of the ligand. Higher RMSDs values were observed for ligand-free structure P1 helix region which affected the helix-join-helix region (C46 to G50) in all the cases. This is clearly indicated by the differences in the superimposed structures of both systems (Figure 3). This result is consistent with the study done by Wood et al. (2012) and Li et al. (2019).

**Table 2:** Average RMSD values of the various structures (complete systems and their different regions).

| Systems | Average RMSD for ligand-free<br>(nm) | Average RMSD for ligand-bound<br>(nm) |
| --- | --- | --- |
| Whole | 0.644 | 0.590 |
| P1 helix | 1.082 | 0.887 |
| P2 helix | 0.278 | 0.284 |
| P3 helix | 0.682 | 0.634 |
| Catalytic pocket | 0.585 | 0.595 |

**Figure 2:**
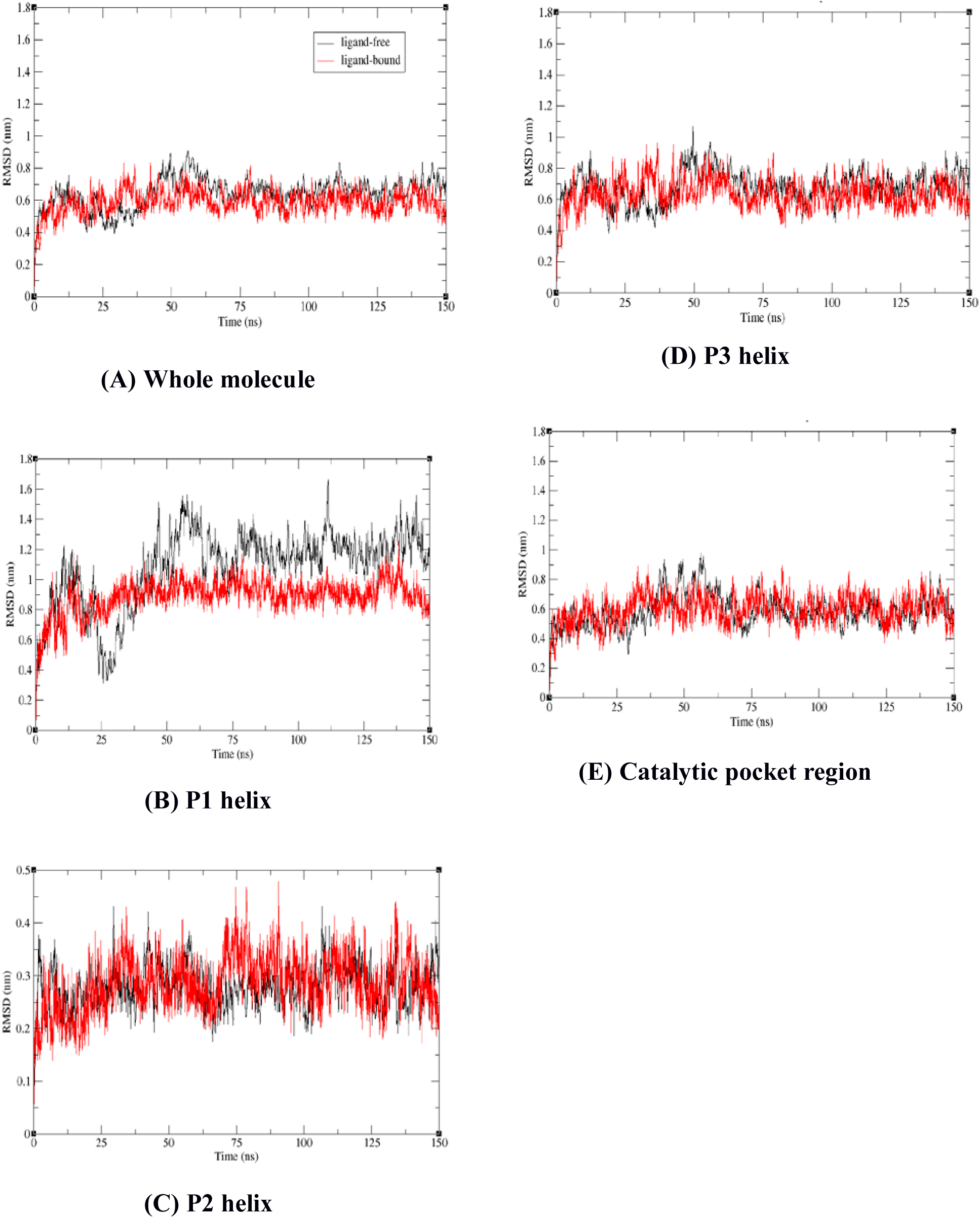
All atoms RMSD plots for ligand-bound and ligand-free systems during 150 ns simulation. **(A)** The graph shows stability in structures after 75 ns and represents complete system RMSD comparison while **(B)** to **(E)** represent RMSD values of the different regions of the c-di-GMP I riboswitch aptamer with respect to the starting minimised structure at the production run. Here the black colour represents ligand-free structure fluctuations and red colour represents ligand-bound structure fluctuations.

**Figure 3:**
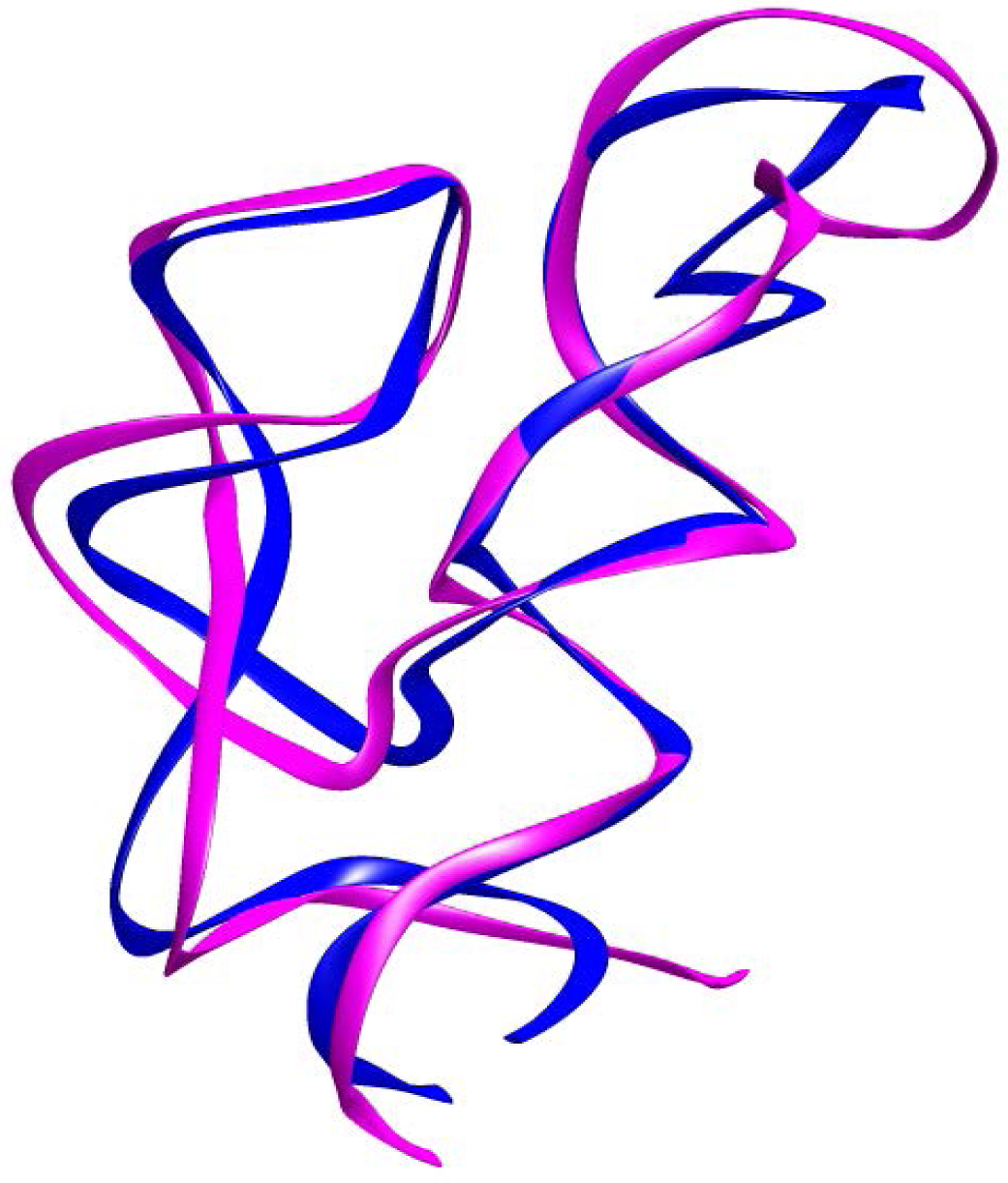
Superimposed structures of the average structures of ligand-free and ligand-bound c-di-GMP I riboswitch aptamer structures. The figure shows changes at P1, P2 and P3 helices. Blue colour represents ligand-bound structure and magenta colour represents ligand-free structure.

### Binding pocket Analysis

The binding pocket key residues and their interactions with the ligand molecule were observed using chimera and gromacs commands. The key residues along with their residue number are reported in Table 3.

**Table 3:** Details of the catalytic pocket generate using Chimera tool.

| S. no. | Nucleotide residue | Nucleotide residue number |
| --- | --- | --- |
| 1 | G | 14 |
| 2 | C | 15 |
| 3 | A | 16 |
| 4 | C | 17 |
| 5 | **A** | **18** |
| 6 | G | 19 |
| 7 | **G** | **20** |
| 8 | G | 21 |
| 9 | **C** | **46** |
| 10 | **A** | **47** |
| 11 | A | 48 |
| 12 | A | 49 |
| 13 | G | 50 |
| 14 | **C** | **92** |
| 15 | C | 93 |

RNA has inherent flexibility and conformational dynamics. In the crystal structure the ligand c-di-GMP (C2E) forms 22 bonds with the surrounding nucleic bases and two crystalline water molecules are present (Figure 4 shows hydrogen bond with surrounding nucleotide residues).

A graph was plotted to observe the number of hydrogen bonds formed between the riboswitch binding pocket residues and the C2E ligand for the entire 150 ns MD run (Figure 5). The average hydrogen bond number is 8.735 (∼9) per frames during the whole simulation time. However, the hydrogen bond number after 110ns shows relatively less hydrogen bonds. The hydrogen bond obtained from the PDB structure using chimera tool showed 12 hydrogen bonds between binding pocket nucleotide residues and the c-di-GMP I ligand. The hydrogen bonds show the strength of binding of ligand to the riboswitch. The residues forming the hydrogen bonds are A18, G20, C46, A47 and C92 (bold faced in Table 3).

**Figure 4:**
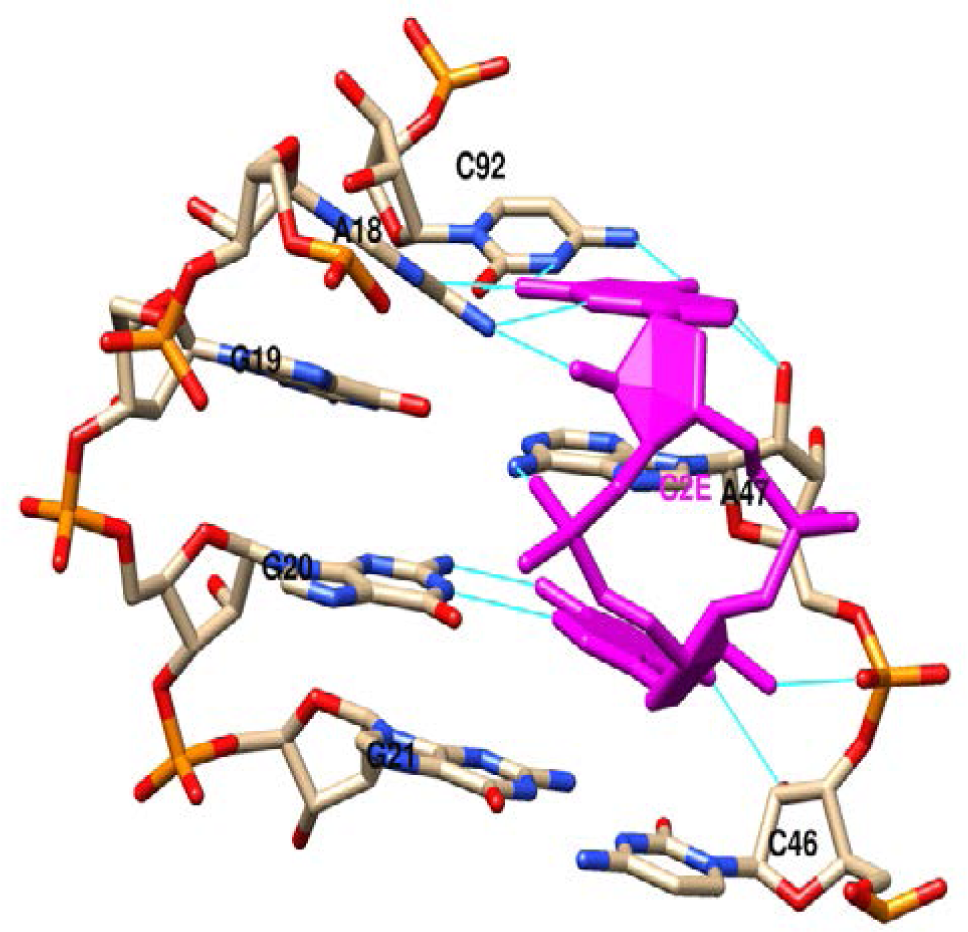
Hydrogen bonding and nucleotide details of the binding pocket of c-di-GMP I riboswitch aptamer. Hydrogen bonds are represented by cyan colour and magenta colour shows the C2E ligand.

**Figure 5:**
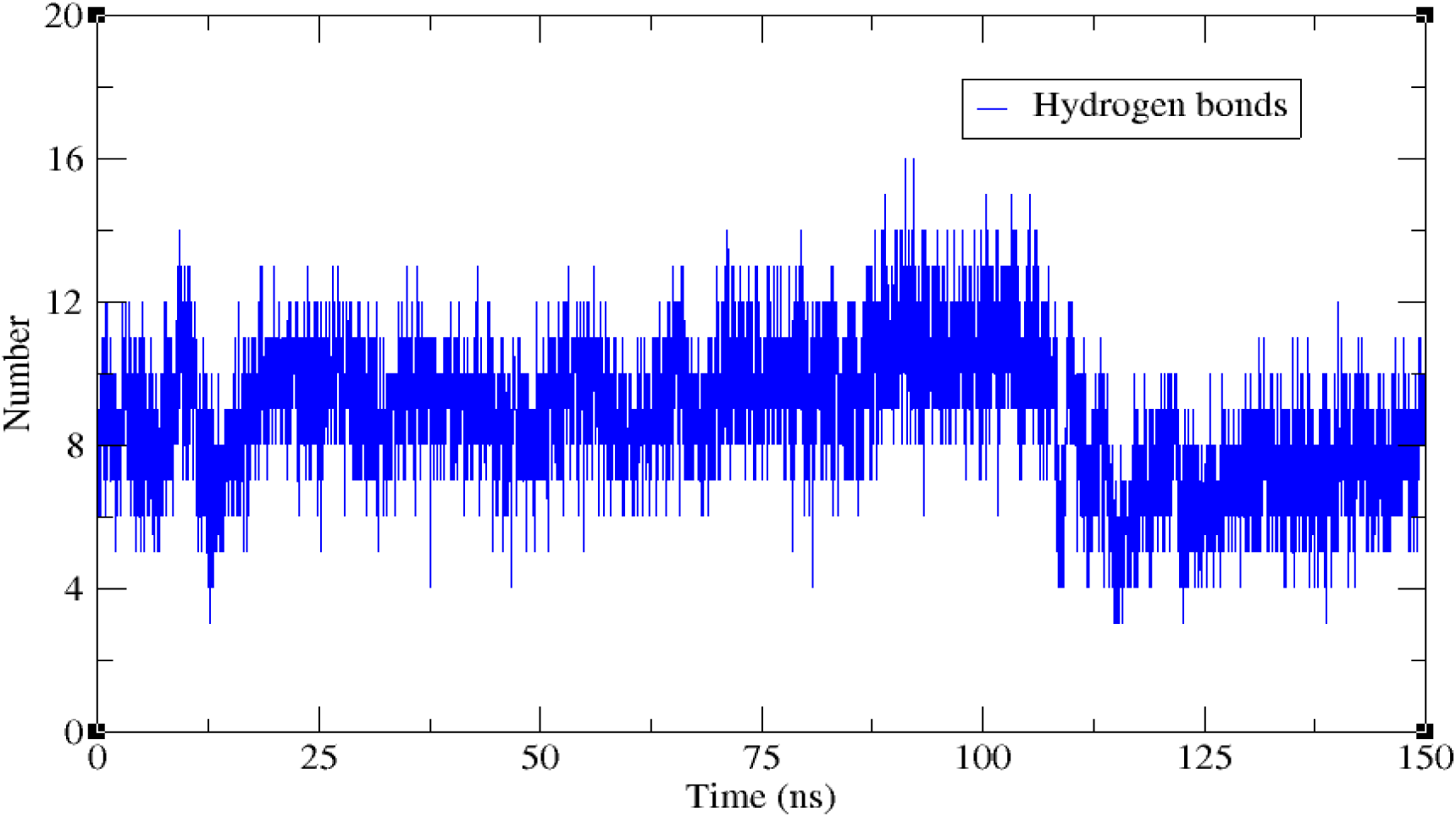
Temporal changes in hydrogen bond numbers between the c-di-GMP I riboswitch aptamer and C2E ligand for 150 ns MD simulation.

The radius of gyration plot for the catalytic pocket clearly illustrated that the binding pocket is not compact rather expanded or open in the ligand-free aptamer, but the binding pocket remained closed for ligand-bound structure (Figure 6).

**Figure 6:**
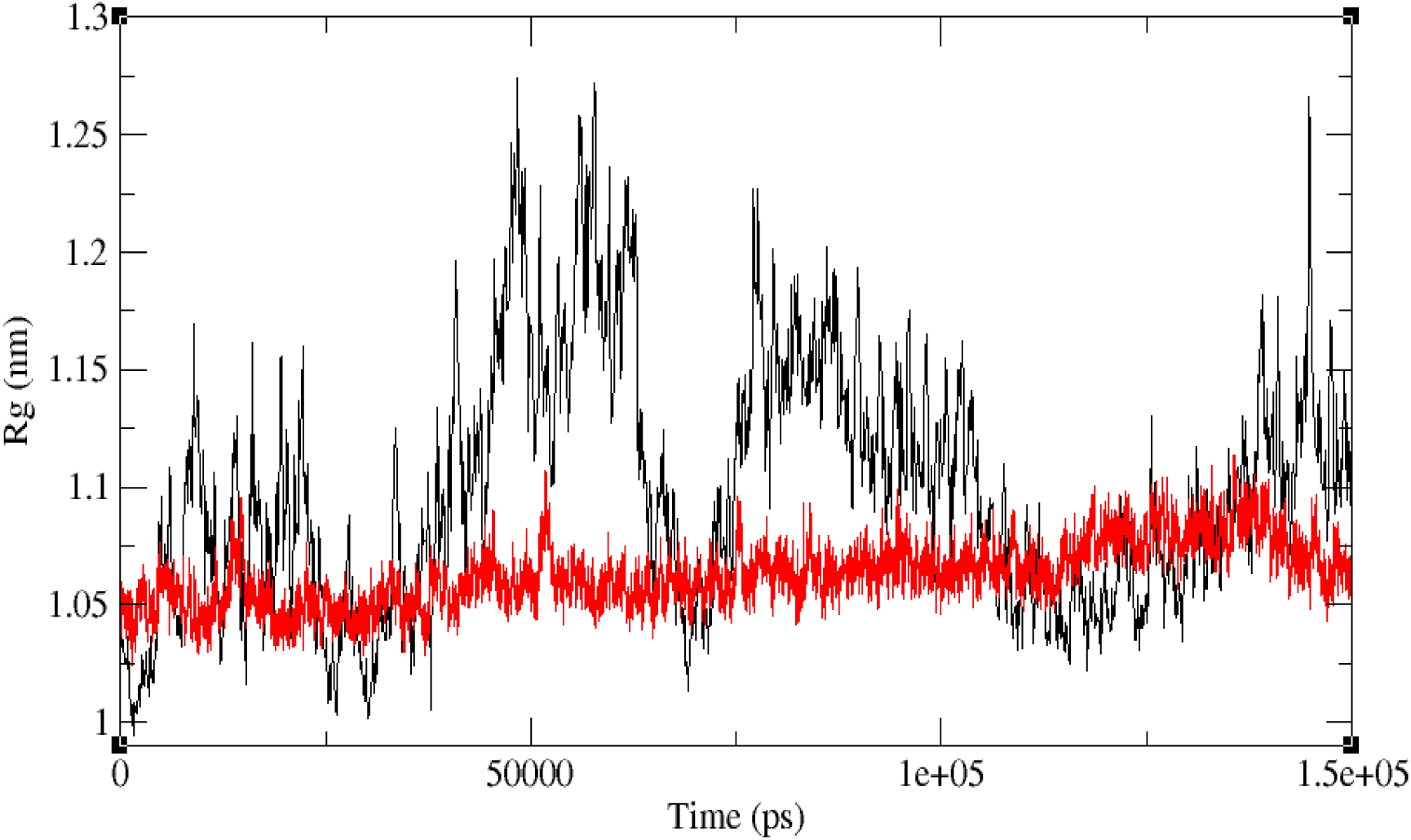
The radius of gyration graph for the binding pocket of ligand-free and ligand-bound aptamer structure.

### Essential dynamics and correlation analysis

To identify the dominant ligand-free motions of the apo_RNA, PCA was applied to the all backbone atoms of the apo_RNA riboswitch. The percentage contribution of the total variance of top twenty eigenvectors is shown in Figure 7 down right, it clearly shows that the top 20 principle components(PCs) captures over 98% of internal motions. It also depicts the first three principal components (PC1, PC2 and PC3) describe about 90% of all the motions. The PC1 has the highest value (76.80%) and strongly dominates the overall variance showing the global dynamics. An assessment of the internal motions through the first three PCs reveals that PC1and PC2 is prominently associated.

**Figure 7:**
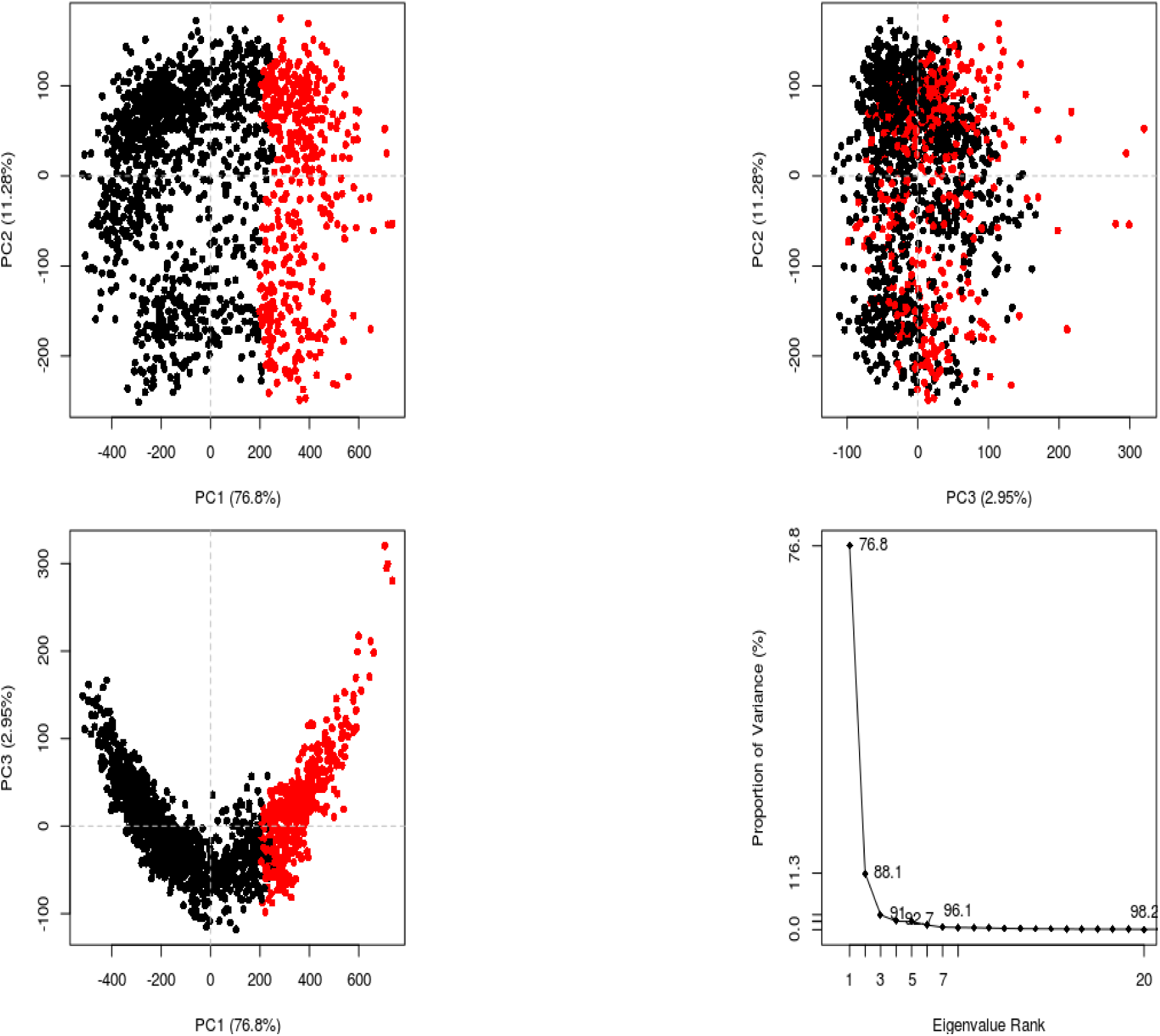
Principle component analysis of apo_RNA MD trajectories. 2D plots for the first three principal components PC1 Vs PC2, PC2 Vs PC3 and PC1 Vs PC3 respectively are represented by first three plots. Here the red and black colour shows two conformational subspaces. The last plot shows the proportion of variance contributed by first 20 PCs which covers 98% internal motions.

Essential dynamics extracts the correlated motions of biomolecules to understand the motions that are most fundamental to the functional activity and partitions conformational spaces into an essential subspace with a few degrees of freedom. The two dimensional plot of PC1 and PC2 shows two major distinct conformational sub-states.

### Dynamic cross correlation (DCC) analysis

DCC are the dynamical changes of the system over the time, it calculate the correlation of the movement of atoms within a molecule that is the degree to which they move together (Kasahara et al, 2014).The DCC map for ligand-free aptamer representing correlation coefficients calculated as time average over duration of simulation is shown in Figure 8. Here the correlation range is from +1 to −1 showing both positive and negative correlation by variations in the colour intensity where blue colour represent positive correlation, pink represent negative correlation and white represent uncorrelated fluctuations. The map revealed that the junction residues are negatively correlated showing pink islands, however the junction J2/3 and J1/3 show correlated movements. The two arms of the P1helix show negative correlated fluctuations illustrating the opposite movement that is drifting apart patterns. The blue islands which indicate positively correlated fluctuations of nucleotides show the formation of the aptamer fold. This supports the existence of a preformed aptamer fold in ligand-free structure.

**Figure 8:**
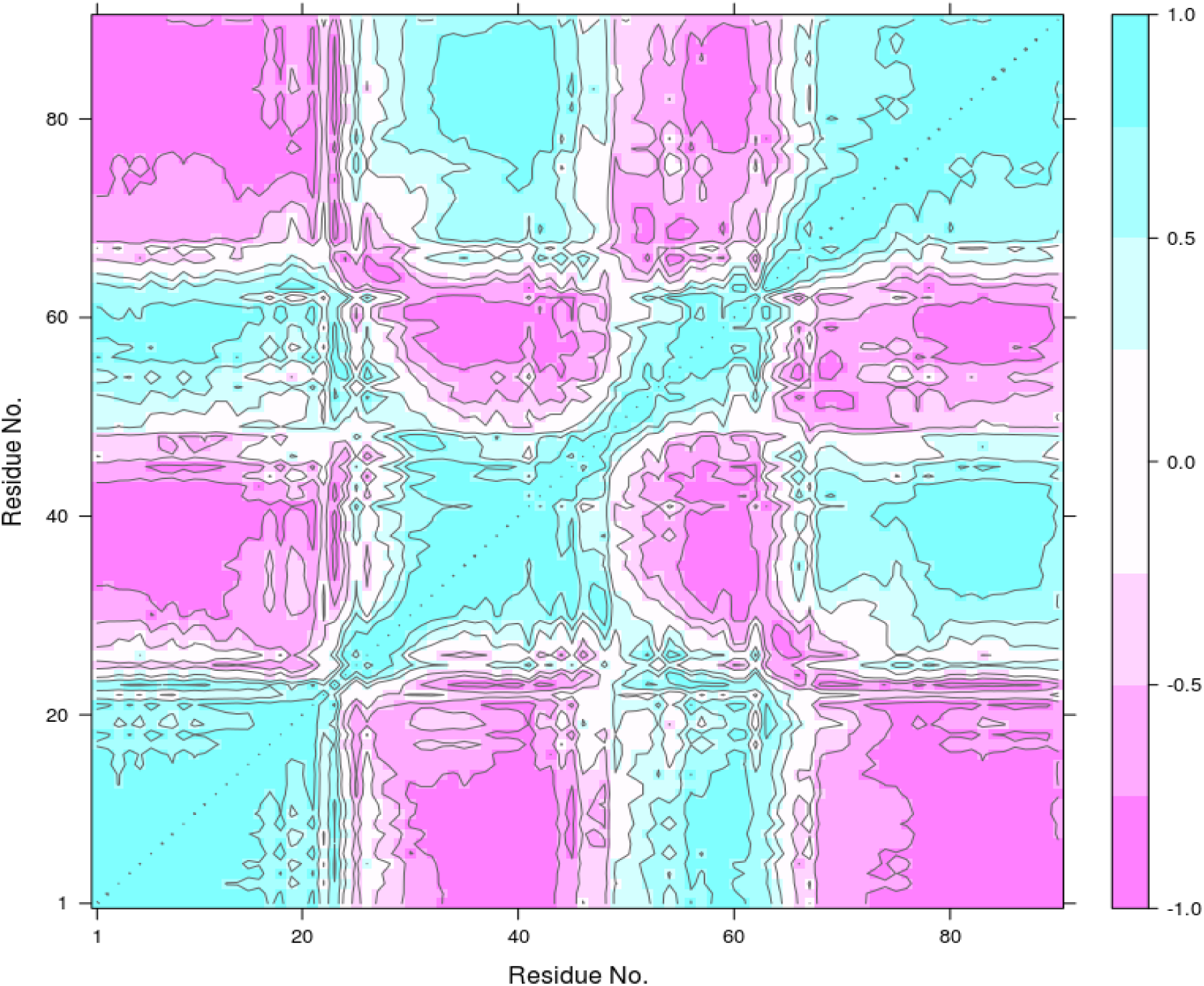
DCC map of 150 ns simulation for ligand-free complex. Blue and pink colour corresponds to high correlations and high anti-correlations respectively, while white regions represent no correlation zone.

Thus our simulations results have shown that the ligand-free structure of the c-di-GMP I riboswitch aptamer have a stable conformation with open binding pocket. This is in agreement with the literature where it had been shown that the ligand-free riboswitch existed in wide open form and have its P2 and P3 stems far apart (Wood et al., 2012). However, in case of ligand-bound structure both the stems are near to each other and form pseudoknot like structure so the junction J1/2, J2/3 and J1/3 are close enough to make bonds with ligand and enclose it in the pocket. Similarly a chemical probing study by Inuzuka et al. (2018), on full length c-di-GMP I riboswitch structure showed translational repression due to masking of ribosome binding site as a result of conformation change in the expression platform on binding of ligand to the aptamer of the riboswitch.

### RMSF analysis

RMSF are the fluctuations of individual atoms of the riboswitches for the simulated time. This provides the insight of motion at atomic level. To understand the dynamical motion further as a comparison between ligand-free and ligand-bound structure, we plotted the RMSF graph for nucleotide as well as atomic fluctuations. The observations from Figure 9 revealed that the fluctuations of nucleotides for ligand-free structure is more than ligand-bound for each nucleotide due to the less stability of ligand-free structure. The ends of aptamer showed higher distance values due to their free motion. A peak at 60-70 nucleotide region show high fluctuations which represent the upper portion of P3 helix in the crystal structure. This area is highly dynamic and flexible as it is free to move in free space while the lower portion of P3 helix formed inter-atomic hydrogen bonds with the helix P2. The binding site region (1200 to 1365) shows higher fluctuations for ligand-free structure as compared to the ligand-bound structure indicates that the nucleotide atoms of this region of the free structure are more dynamic and free to move. The smaller peaks at 25, 46, 53 and 62 for ligand-bound structure show slightly higher fluctuations then their corresponding nucleotides or atoms in ligand-bound structure. The residue at 25 correspond to apex of P2 helix which is free to move as it donot participate in forming pseudoknot with P3helix, similarly residues 46, 53 and 62 belong to P3 helix which have free movement due to no hydrogen bond formation with other nucleotide residue.

**Figure 9:**
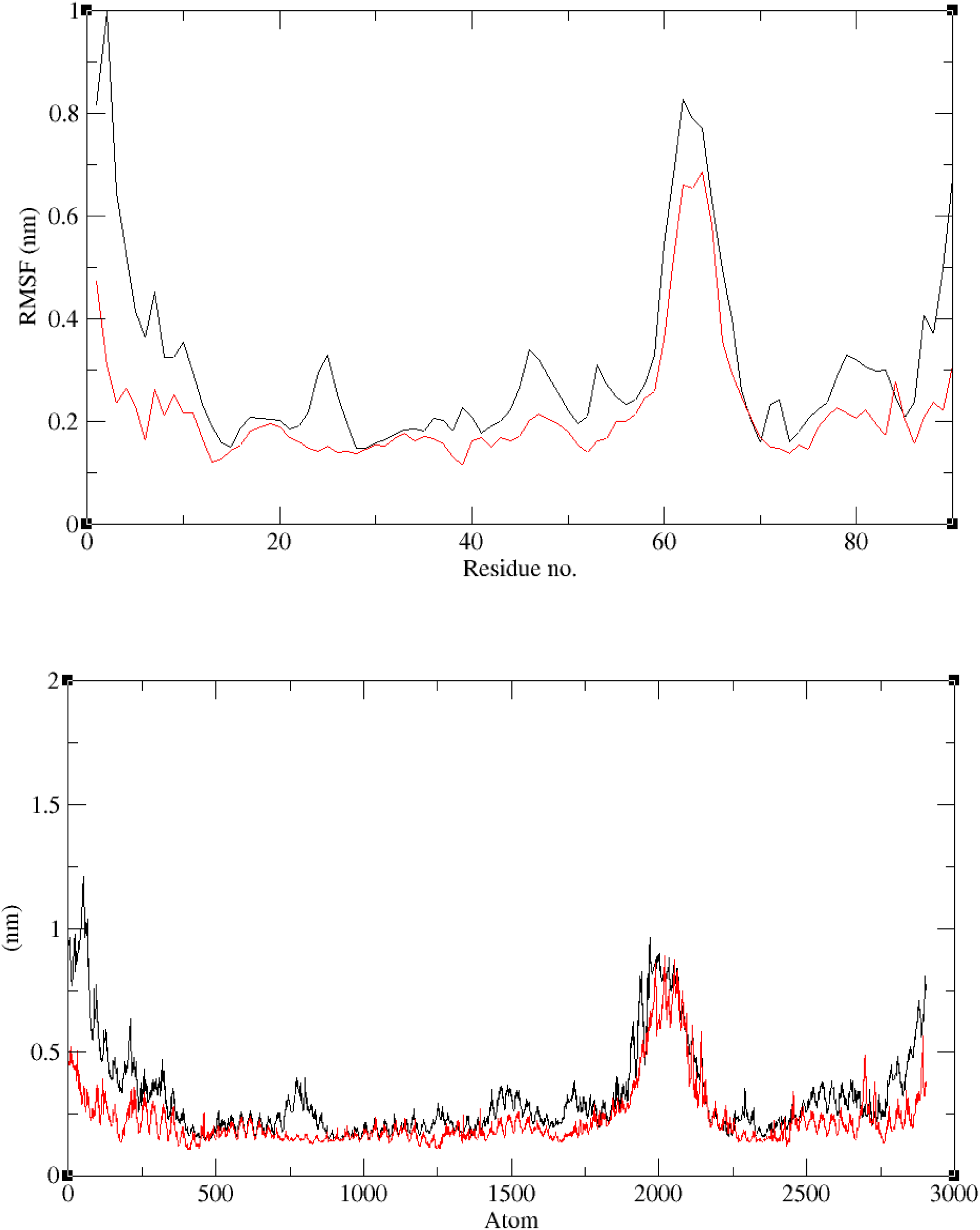
RMSF plots of ligand-bound and ligand-free systems during 150 ns simulation for both atom and nucleotide residue. These plots were generated for the RNA molecule of both the systems and the reference structure used is the initial structure during MD production run. The peak region represents P3 helix secondary structure.

### SASA analysis

SASA is the calculation of the surface area of the biomolecule (protein or RNA) interacting with water or the area exposed to water. The graph shows a difference (6.09 nm^2^) in the average surface area of the fluctuations as a function of time for both ligand-free and ligand-bound structures (Figure 10). This difference might be due to the expansion of ligand-free structure as stated by the radius of gyration plot earlier. Further the SASA graph was plotted for the binding pocket which revealed that the ligand-free structure showed relatively more solvent accessible surface area till 110 ns and then SASA for ligand-bound structure increased (Figure 10 A). The average SASA difference between ligand-free and ligand-bound structures for binding pocket is 0.27 nm^2^, this minor difference may be due to exposure of part of A47 residue which gets capped due to the U shape structure of ligand molecule.

**Figure 10:**
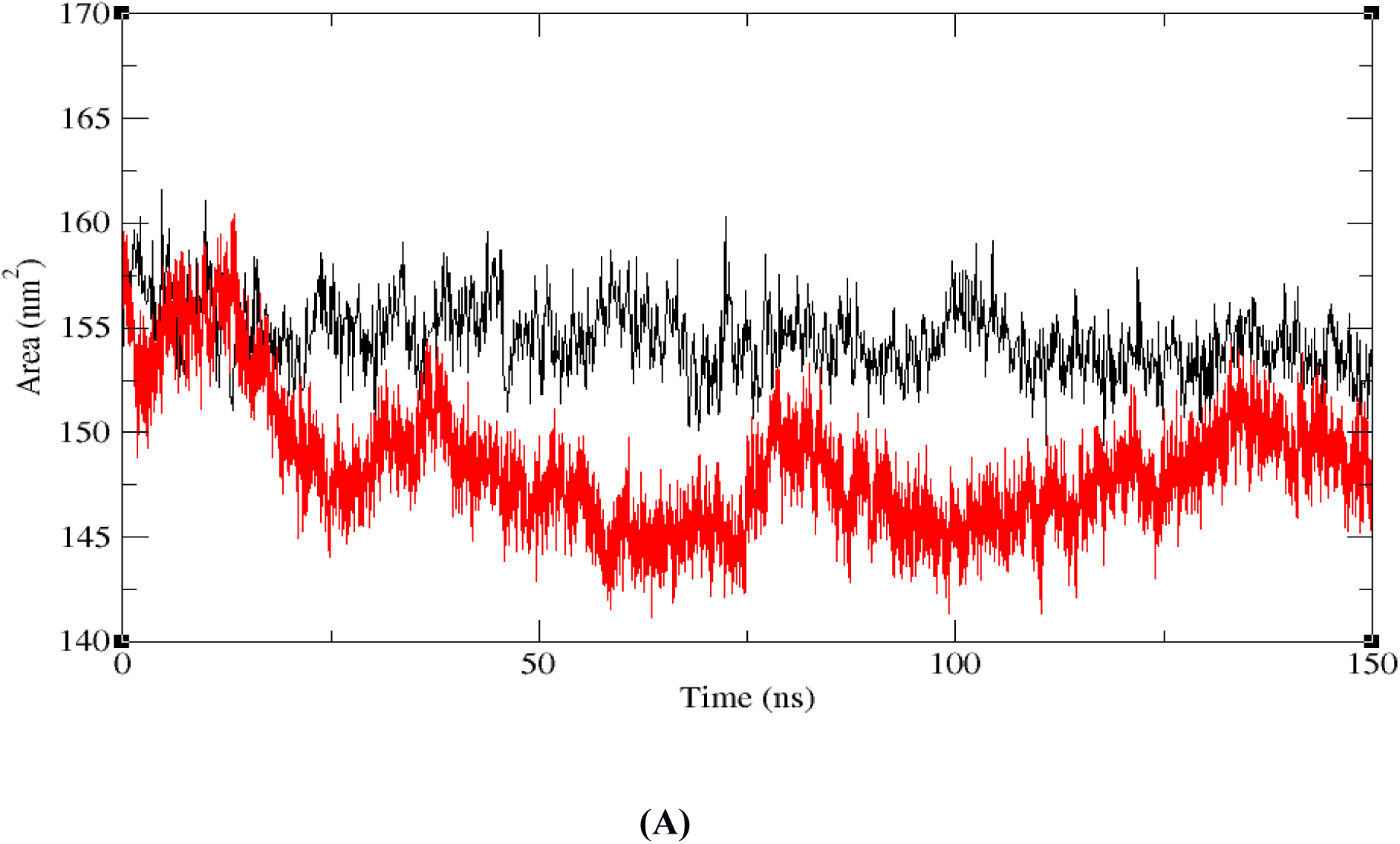
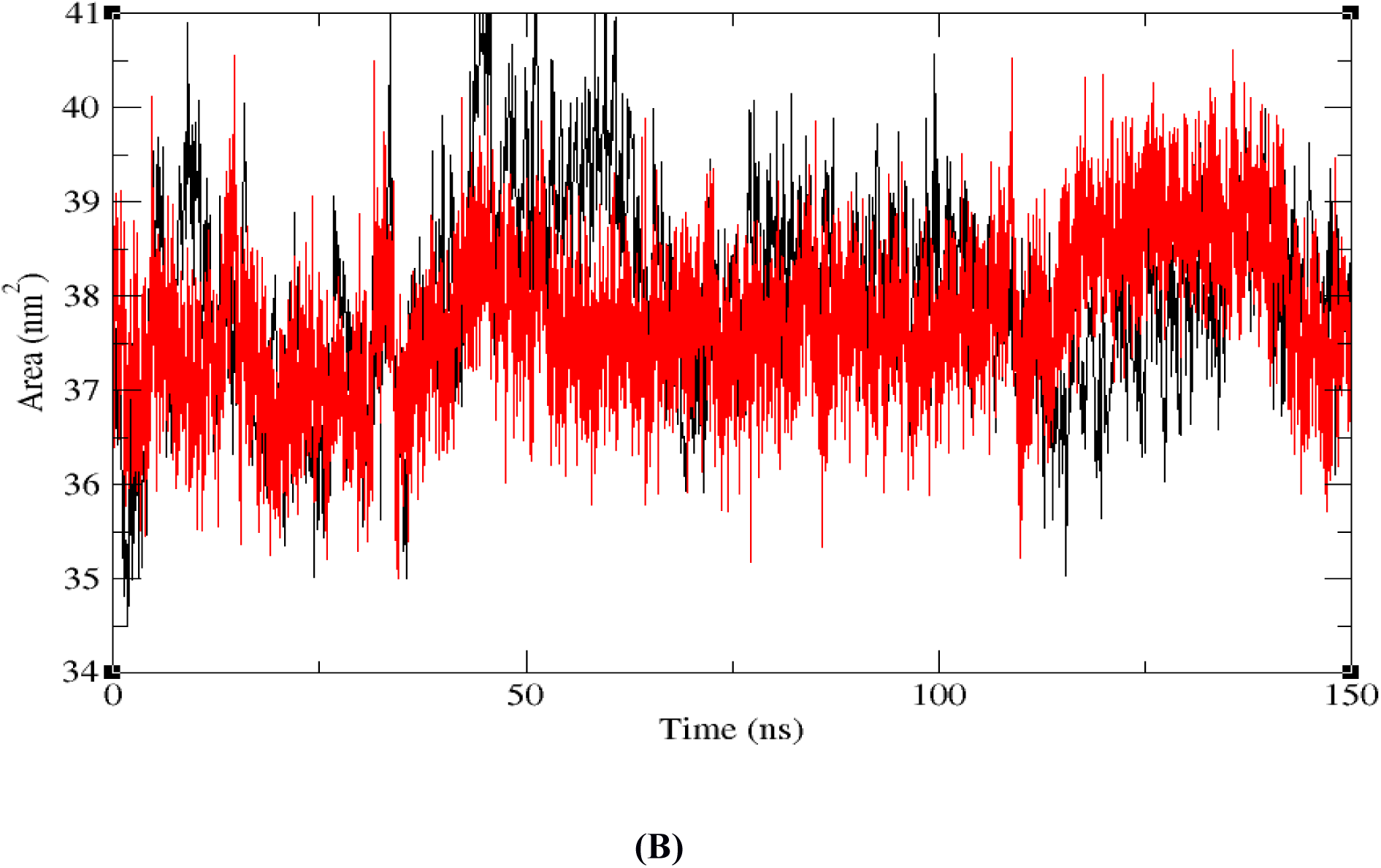
SASA plots for ligand-bound and ligand-free systems during 150 ns simulation. The graph **(B)** represents SASA calculations for the binding pocket nucleotides.

## Conclusions

MD simulations had been performed for both the ligand-bound and ligand-free aptamer domain of the c-di-GMP I riboswitch to identify ligand-free stable structure and the role of aptamer in recognition of specific ligand and binding mechanism. The simulations resulted in identification of ligand-free aptamer structure with stable conformation which has folded P2, P3, open catalytic pocket and an unwind P1 helix. The results also showed that the binding of ligand to the aptamer plays an important role in stabilizing the overall structure. In the absence of c-di-GMP I, the aptamer showed changes in the binding pocket region showing a shift in the residues A18, G20, C46, A47 and C92. These represent the key residues which play critical role in ligand binding and are found to be strongly conserved in the bacterial system. Higher stable fluctuations were seen at the P1 helix region while the P2 and P3 helices remained less dynamic as compared to the trajectories dynamics of the ligand-bound structure. Thus the trajectory and correlation analyses demonstrated the interactions in the secondary structures to illustrate the binding mechanism. These results might be helpful to understand conformational interactions between the aptamer and expression platform of c-di-GMP I riboswitch further. The presented study is useful in finding high affinity inhibitors for c-di-GMP I riboswitch which act as potential RNA drug target. Further, this study can be expanded to study mutations in the key residues and their effect on the binding affinity.

## Abbreviations

ACPYPE: AnteChamber PYthon Parser interfacE
AMBER: Assisted Model Building with Energy Refinement
C2E: Cyclic Diguanosine Monophosphate
C-di-GMP I: Cyclic Diguanosine Monophosphate I
DCC: Dynamic Cross Correlation
GROMACS: GROningen MAchine for Chemical Simulations
MD: Molecular Dynamics
mRNA: Messenger Ribonuceic acid
NPT: Number of atoms, Pressure and Temperature
ns: nano second
NVT: Number of atoms, Volume and Temperature
PDB: Protein Data Bank
PME: Particle Mesh Ewald method
RMSD: Root Mean Square Deviation
RMSF: Root Mean Square Fluctuations
SASA: Solvent Accessible Surface Area
UTR: Untranslated regions
VMD: Visual Molecular Dynamics

## Authors’ contribution

Conceived, designed and performed the experiments: PK. Analyzed the data and wrote the paper: PK and AS. Supervised the research: AS.

## Acknowledgement

We thank Dr. Dhananjay Bhattacharya, Dr. Jens Carlsson and Dr. Ramasubbu Sankararamakrishnan for various useful suggestions. PK thanks Dr. Mark E Tuckerman and Dr. Nishant Nair for useful discussions. AS thanks DBT, India for financial support. PK thanks the UGC, India for providing financial assistant to carry out research work.

